# Sparsely-Connected Autoencoder (SCA) for single cell RNAseq data mining

**DOI:** 10.1101/2020.05.26.117705

**Authors:** L Alessandri, F Cordero, M Beccuti, N Licheri, M Arigoni, M Olivero, F Di Renzo, A Sapino, RA Calogero

## Abstract

Single-cell RNA sequencing (scRNAseq) is an essential tool to investigate cellular heterogeneity. Although scRNAseq has some technical challenges, it would be of great interest being able to disclose biological information out of cell subpopulations, which can be defined by cluster analysis of scRNAseq data. In this manuscript, we evaluated the efficacy of sparsely-connected autoencoder (SCA) as tool for the functional mining of single cells clusters. We show that SCA can be uses as tool to uncover hidden features associated to scRNAseq data. Our approach is strengthened by two metrics, QCF and QCM, which respectively allow to evaluate the ability of SCA to reconstruct a cells cluster and to evaluate the overall quality of the neural network model. Our data indicate that SCA encoded spaces, derived by different experimentally validated data (TFs targets, miRNAs targets, Kinases targets, and cancer-related immune signatures), can be used to grasp single cell cluster-specific functional features. In our implementation, SCA efficacy comes from its ability to reconstruct only specific clusters, thus indicating only those clusters where the SCA encoding space is a key element for cells aggregation. SCA analysis is implemented as module in rCASC framework and it is supported by a GUI to simplify it usage for biologists and medical personnel.

## Introduction

Single-cell RNA sequencing (scRNAseq) has emerged as essential tool to investigate cellular heterogeneity. Single cell analysis is instrumental to understand the functional differences existing among cells within a tissue. Individual cells of the same phenotype are commonly viewed as identical functional units of a tissue or organ. However, single cells sequencing results [1] suggest the presence of a complex organization of heterogeneous cell states producing together system-level functionalities. Network analysis is a crucial tool to uncover biological and pathological mechanisms, and it is becoming an area of research for the single cell bioinformatics community. Recently, Pratapa and colleagues [2] benchmarked 12 gene networks tools for scRNAseq, notably none of these tools exploits neural network approaches to discover functional features associated with cells’ clusters. A particular type of neural network, autoencoder, seems to be particularly interesting and suitable for the analysis of single cell data. Autoencoder is an unsupervised artificial neural network, which efficiently compresses and encode data, then learns how to reconstruct the data back from this reduced encoding to produce an output that is as close as possible to the original input [3]. Autoencoder reduces data dimensions by learning how to ignore the noise in the data. Autoencoder-based approaches have been used to cluster single cell data [4], to impute single cell data [5], data denoising [6], batch correction [7]. Recently, Gold and co-workers [8] have evaluated the use of autoencoders for data interpretation, implementing sparsely-connected autoencoder (SCA) to gene set analysis. SCA uses a single-layer autoencoder with sparse connections (representing known biological relationships) in order to attain a value for each gene set. SCA provides great flexibility for modelling biological phenomena [8].

We made available to the single-cell community a framework, rCASC [9], providing an integrated analysis environment for single-cell RNAseq. rCASC provides all the tools for sub-population discovery, which can be achieved using different clustering techniques, based on different distance metrics [9]. In this manuscript, we introduce a new rCASC module for functional annotation of cell clusters using SCA.

## Results

### Sparsely-Connected Autoencoder (SCA)

SCA encoding/decoding functions consist of a single sparse layer (Fig. 1, latent space), with connections based on known biological relationships [8, 10]. Each node represents a known biological relationship (transcription factors (TFs) targets, miRNA targets (miRNAs), cancer-related immune-signatures (ISs), kinase specific protein targets (Ks), etc.) and only receives inputs from gene nodes associated with the biological relationship. With respect to the Gold paper [8], which uses gene sets [11], in our implementation the latent space is based on experimentally validated data, TRRUST [12], miRTarBase [13], RegPhos [14], and a manually curated cancer-based immune-signature (See material and methods).

**Fig. 1:**
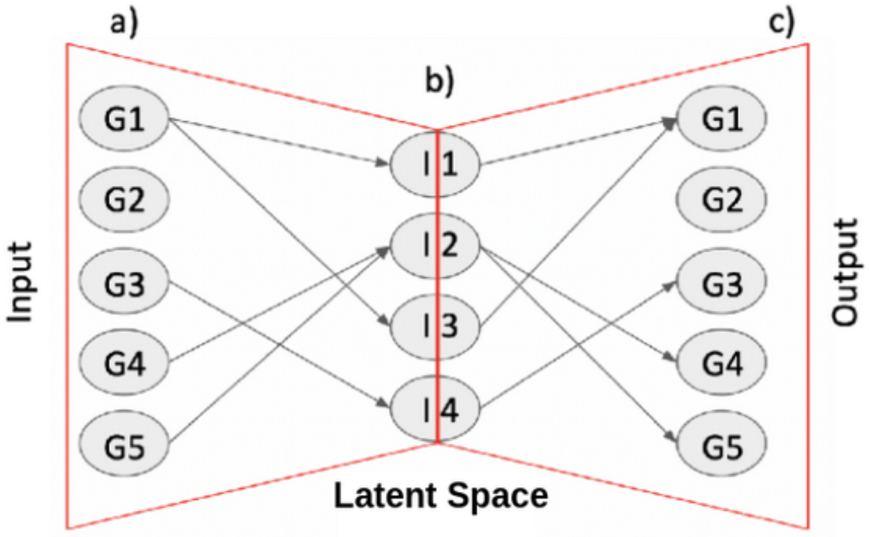
Shallow Sparsely-Connected Autoencoder (SCA) architecture. a) gene-level expression profile for each cell are used as input. b) The hidden layer is made of nodes where each node is a transcription factor, a miRNA or a functional signature. The vertices connecting input/output (c) nodes are based on experimentally validated biological knowledge.

SCA analysis is executed multiple times on a cell dataset, previously partitioned in clusters, using any of the clustering tools implemented in rCASC: tSne+k-mean [15], SIMLR [16], griph [17], scanpy [18] and SHARP [19]. Cell Stability Score (CSS) [9] is used on SCA outputs to generate two quality scores metrics: QCF and QCM. QCF is generated comparing the cell partition generated by rCASC clustering, i.e. the reference clusters, with the clusters generated in each SCA run. This metric measures the efficacy of the latent space in describing cells aggregations corresponding to at least part of the reference clusters. A SCA cluster is considered comparable to the corresponding reference cluster if QCF mean is ≥ 0.5, where 0.5 indicates that at least 50% of SCA runs retain the structure of the reference cluster. QCM instead measures the consistency of the clusters generated in each SCA run. If a set of biological information is important for the definition of a cluster then the majority of the SCA runs will converge to a similar solution. Thus, comparison, via CSS, of random couples of clusters selected over multiple SCA runs must show a conserved cluster(s) organization. On the basis of our experiments we decided to consider sufficiently robust for further analysis only clusters supported by QCM mean ≥ 0.5, where 0.5 indicated that at least 50% of the compared latent space models retain the same cluster(s) structure. Thus, a “robust cluster”, i.e. a SCA cluster retaining the cell organization of at least one of the reference clusters, should have both QCF and QCM ≥ 0.5.

A frequency matrix for the latent space representations is also built and it is used as input for COMET [20], which is a software able to identify marker signatures specific of each cluster. Only clusters characterized by QCM and QCF means ≥ 0.5 should be used for marker signature detection.

### SCA analysis on a PBMC derived dataset (setA)

A data set (setA), based on FACS purified cell types [21] was used to investigate the SCA behaviour. SetA was previously used to estimate the strength of CSS metric [9]. We clustered setA using all the clustering tools actually implemented in rCASC: tSne+k-mean [15], SIMLR [16], griph [17], Seurat [22], scanpy [18] and SHARP [19]. All tools but tSne+k-mean and scanpy provided very good and similar partition of the different cell types (Supplementary Fig. 1). As clustering tool for testing the behaviour of SCA we used griph [17], Fig. 2A.

**Fig. 2:**
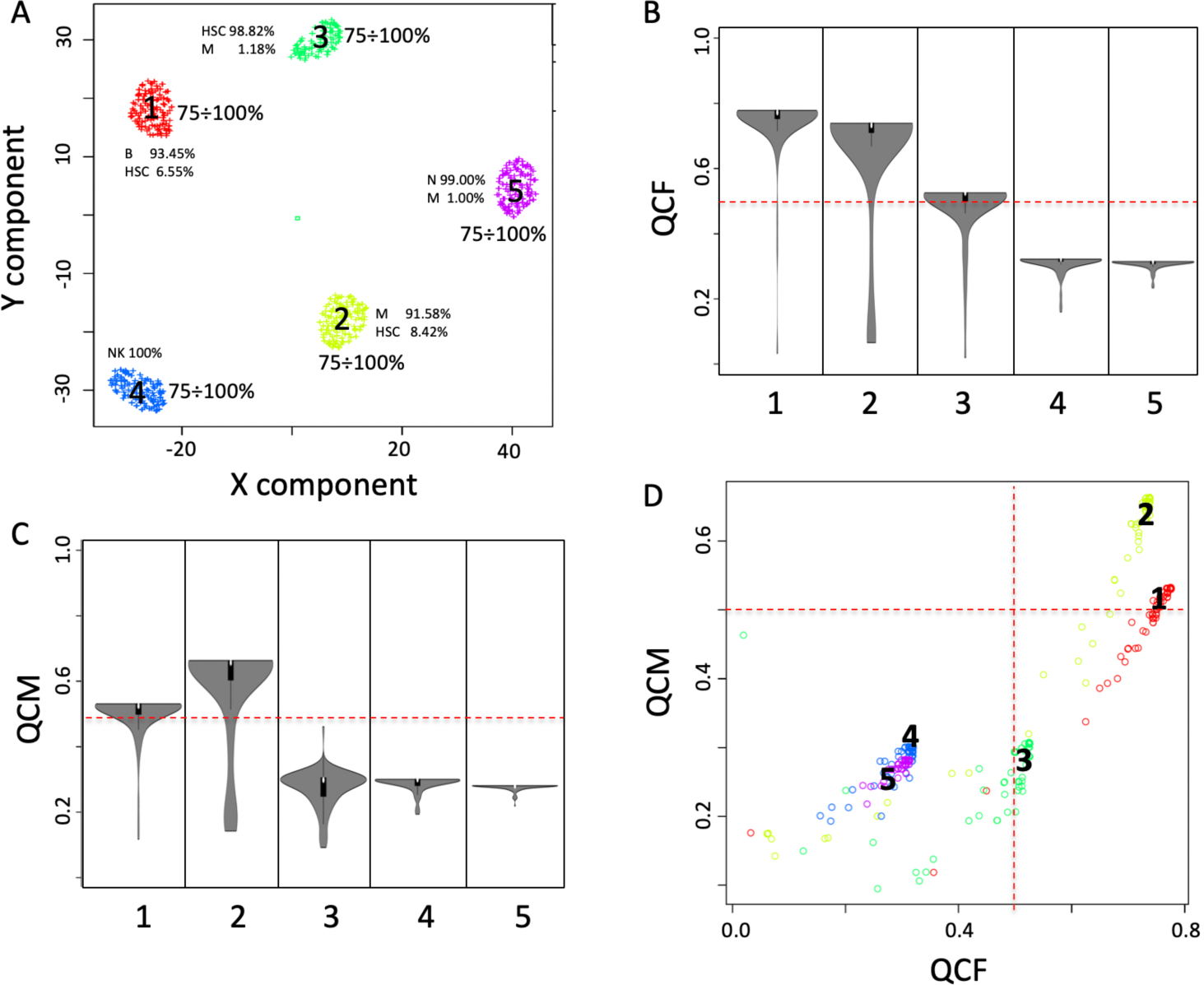
SCA analysis using TFs. A) five clusters were detected analysing setA with griph [17]. Each cluster is made by more than 90% by one cell type. A little amount of HSC is contaminating B cells, monocytes and naïve T cells. B) QCF violin plot. The metric is an extension of CSS and it measures the ability of latent space to keep aggregated cells belonging to predefined clusters (A). The metrics has the range between 0 and 1, where 1 indicates a high coexistence of cells within the same cluster in the bootstrap analysis, and 0 a total lack of coexistence of cells within the same cluster. In this example, only clusters 1 and 2 can be well explained by TF-based latent space. C) QCM violin plot, this metric is also an extension of CSS and it measures the ability of the neural network to generate consistent data over bootstraps. In this example, SCA provides consistent data only for clusters 1 and 2. D) Combined view of QCM and QCF, informative clusters are those characterised by high QCM and QCF scores, in this example only clusters 1 and 2 are characterized by a robust neural network able to keep the cell aggregated using TF-based knowledge. Dashed red line indicates the defined threshold to consider the latent space information suitable to support cells’ clusters.

SCA analysis was done using a TFs-based latent space, where each hidden node is a TF and arches connecting each hidden node represent experimentally validated TF target genes from TRRUST database [12]. From this analysis we observed that only cluster 1 and 2 (Fig. 2A) could be explained by a SCA based on TFs, since only these two clusters are supported by a QCF and QCM ≥ 0.5 (Fig. 2B, C, D).

The matrix describing the frequency of the latent space variables is used to extract cluster specific signatures using COMET tool [20], which is also implemented in rCASC. COMET is a computational framework for the identification of candidate marker panels consisting of one or more genes for cell populations of interest identified with single cell RNAseq data. We analyzed with COMET the frequency matrix of TFs derived from SCA latent space. The four top ranked markers for cluster 1 are PAX5, NFAT5, RFXANK and CHD4, (Table 1, Fig. 3A).

**Table 1.**
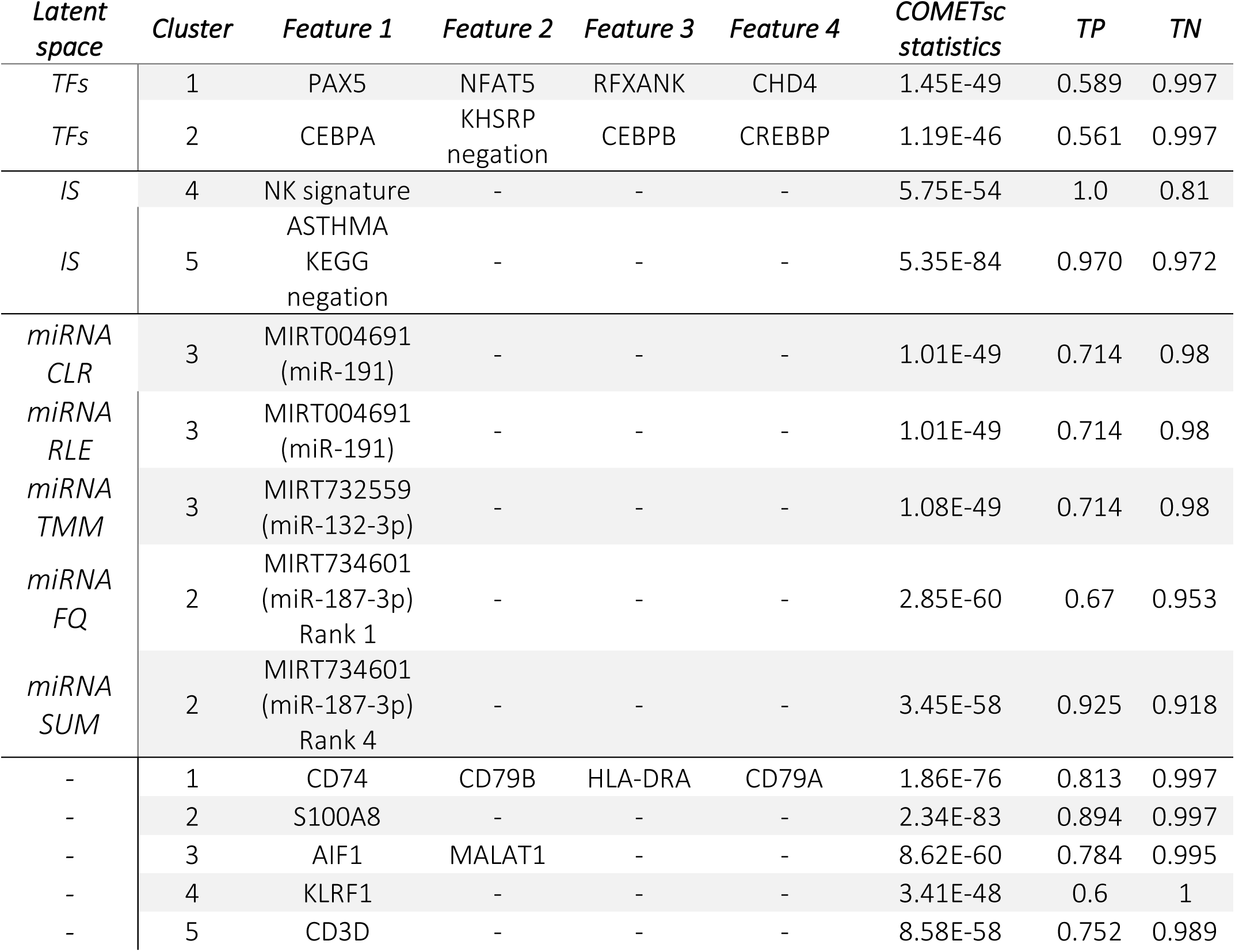

**Fig. 3:**
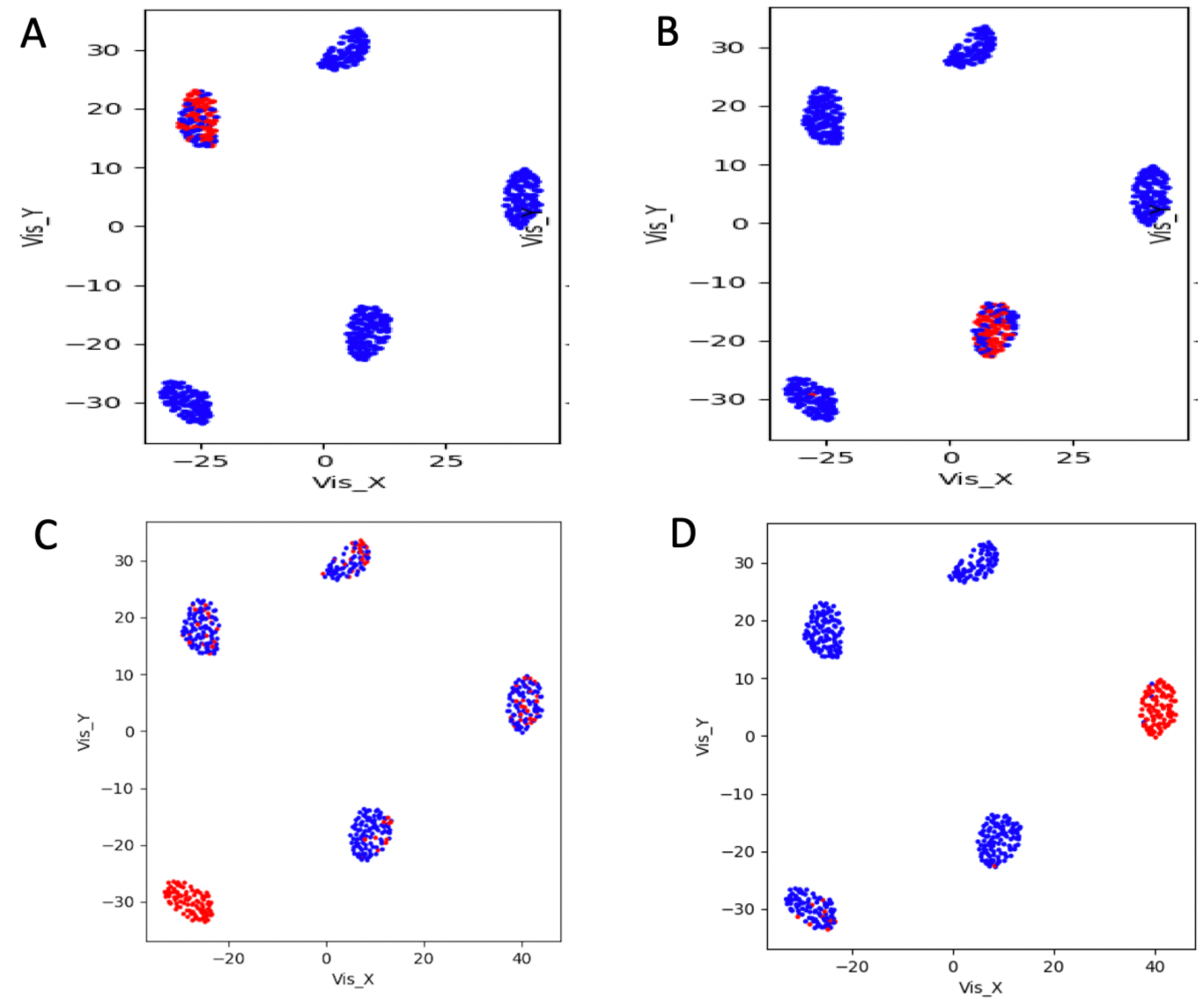
COMET analysis of the latent space matrices. A) Set of 4 genes (PAX5, NFAT5, RFXANK and CHD4) characterizing cluster 1 (B-cells). B) Set of 4 genes (CEBPA, KHSRP negation, CEBPB, CREBBP) characterizing cluster 2 (Monocytes). Blue dots indicate the lack of the 4 markers, red dots the presence of the 4 markers. C) Rank 1 NK signature (30266715) specifically characterizing cluster 4 (NK). D) Rank 2 Asthma KEGG negation, specifically characterizing Naïve T-cells.

The four markers detected for the cluster 1, made mainly by B-cells, recapitulate very well some of the main key elements of this cell type. The transcription factor PAX5 is essential for the commitment of lymphoid progenitors to B-lymphocyte lineage. PAX5 fulfills a dual role by repressing B lineage ‘inappropriate’ genes and simultaneously activating B lineage-specific genes [23]. NFAT5 is essential for optimal antibody productivity [24]. CHD4 is essential for early B-cell development [25] and RFXANK is involved in activation of MHC-II genes. MHC-II molecules are largely restricted to thymic epithelial cells and professional antigen-presenting cells, including dendritic cells, macrophages, and B-cells. Moreover, for cluster 2, mainly composed of monocytes, the four top ranked markers (CEBPA, KHSRP negation, CEBPB, CREBBP, Table 1, Fig. 3B) are strongly involved in monocyte functionalities [26]. The transcription factor CCAAT/enhancer-binding protein β (CEBPB) is highly expressed in monocytes/macrophages and it is a critical factor for Ly6C-monocyte survival [27]. The downregulated expression of the KH-Type Splicing Regulatory Protein (KSRP) during monocytopoiesis and its up-regulated expression during granulopoiesis suggests that KSRP has divergent roles during monocytic and granulocytic differentiation [28]. CREB induces an anti-apoptotic survival signal in monocytes and macrophages [29] and CREBBP specifically binds to the active phosphorylated form of CREB [30].

A SCA based on a manually curated cancer-immune-signature (IS, Fig. 4) was also used. This SCA shows good QCM/QCF for cluster 4 (Natural Killer cells) and cluster 5 (Naïve T-cells).

**Fig. 4:**
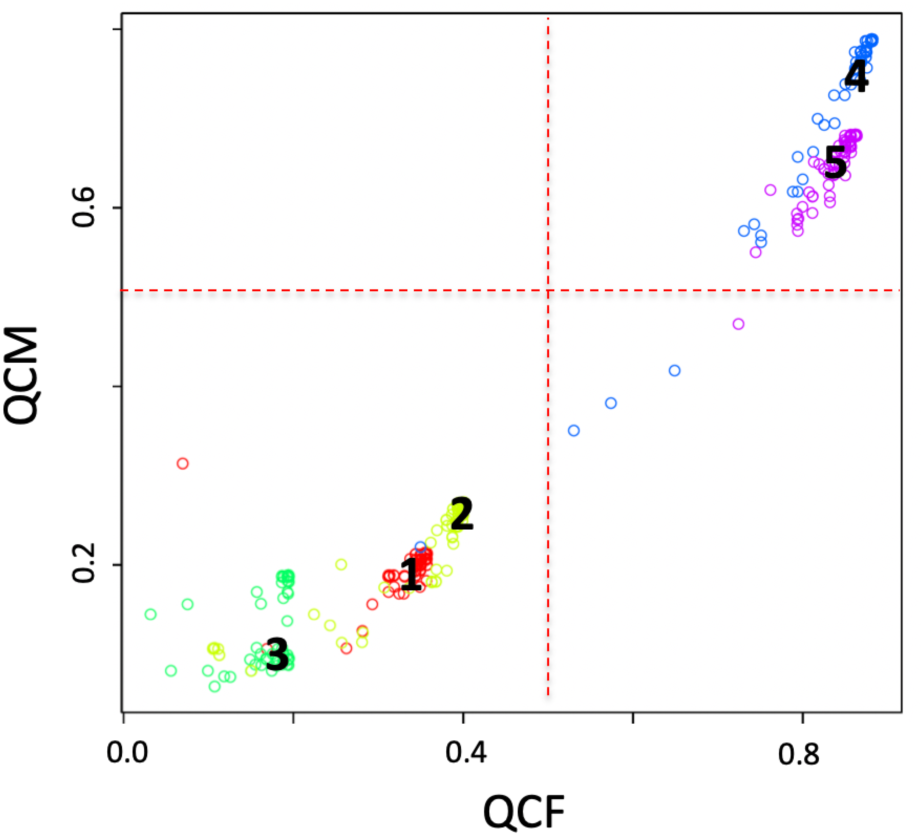
SCA analysis using a manually curated cancer immune-signature. Input counts table for SCA are log10 transformed.

Analyzing the latent space IS frequency table with COMET we identified: one feature of the immune-signature associated to NK (Table 1, Fig. 3C) and a KEGG signature, which negation is specific for Naïve T-cells cluster (Table 1, Fig. 3D). SCA analysis on setA was also done using a latent space based on kinase targets, but we cannot find any robust association with clustering data (not shown).

COMET analysis of the 5 clusters generated by griph analysis on the raw counts table, i.e. reference clusters, detected gene markers (Table 1) that recapitulate cluster specific cell type characteristics: CD79A and B (cluster 1, B-cells) in association with surface Ig constitute the B-cell antigen receptor complex and play a critical role in B-cell maturation and activation. CD74 (cluster 1, B-cells) is an integral membrane protein that functions as MHC class II chaperone present on B-cells. S100A8 (cluster 2, Monocytes) is released by monocytes during inflammation [31]. AIF1 (cluster 3, Hematopoietic Stem Cells) is a main driver of hematopoietic stem cells differentiation [32]. KLRF1 (cluster 4, Natural Killer) is part of NK gene complex on chromosome 12p12.3 [32]. CD3D (cluster 5, naïve T cells) is a specific receptor of T-cells. Notable, none of the TFs detected by SCA analysis are present in the COMET results generated using the raw counts table.

### Effect of normalization on SCA latent space frequency matrix

SCA based on validated target genes for miRNAs (Fig. 5A) shows that clusters 3 and 4 have a potentially interesting trend, although they are not supported by a QCM and a QCF ≥ 0.5. Thus, we have investigated the effect of various normalization procedures of the SCA input counts table on the modelled results. In Fig. 5, it is clear that normalization affects both QCM and QCF for clusters which are below but close to the suggested threshold (0.5), i.e. cluster 3 (Fig. 5A, B, E) cluster 2 (Fig. 5C, D). Low quality clusters, i.e. those in which there is clearly no information that is gathered from SCA latent space are minimally affected by normalization, i.e. clusters 1, 4 and 5. This observation suggests that it is important to assess the effect of different normalization procedures on SCA input data, specifically if clusters are located near to the significant threshold suggested for QCM and QCF.

**Fig. 5:**
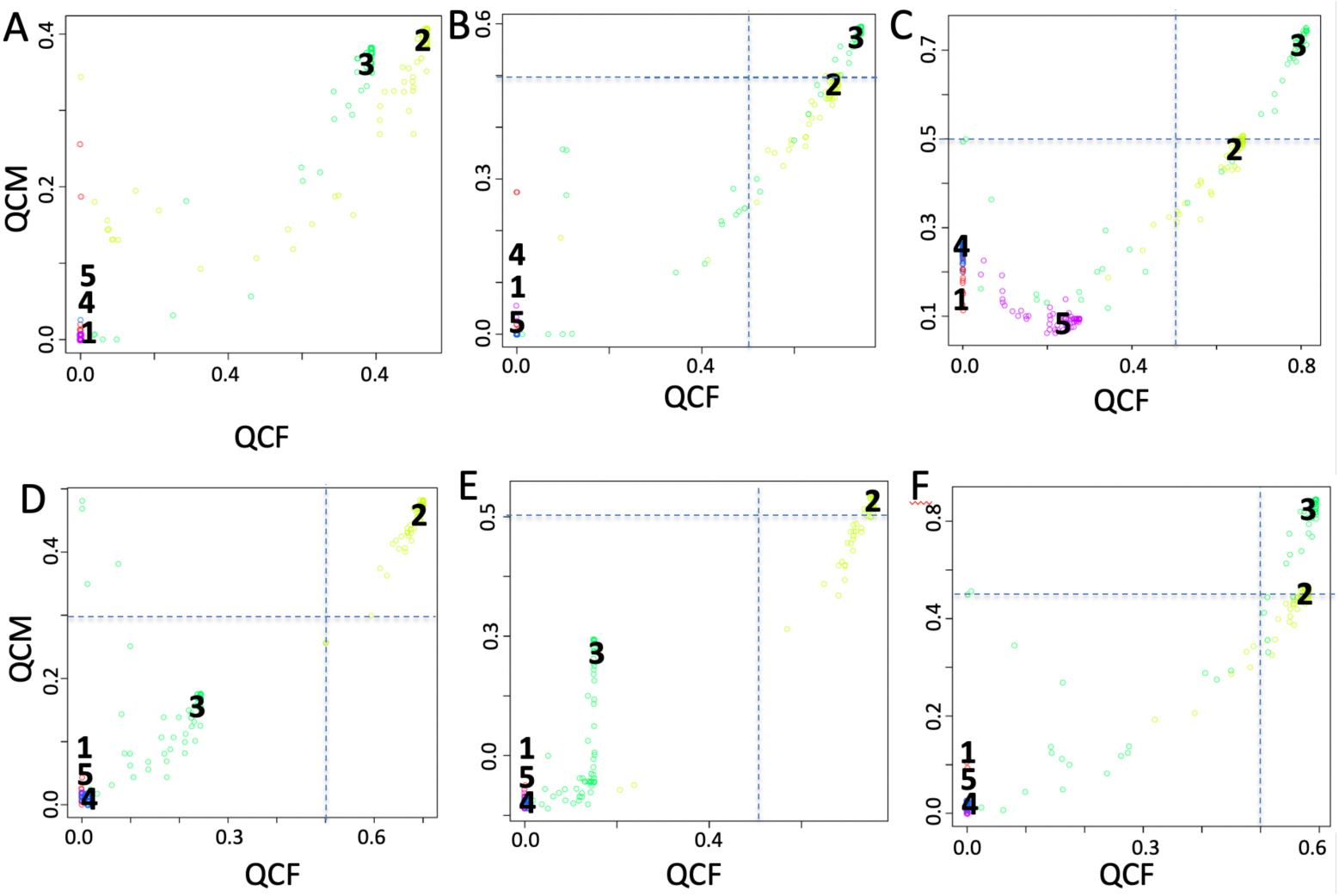
Effect on QCM/QCF as a consequence of the normalization of the SCA input counts table. A) Log10 transformed, B) Centred log-ratio normalization (CLR), C) relative log-expression (RLE), D) full-quantile normalization (FQ), E) sum scaling normalization (SUM), F) weighted trimmed mean of M-values (TMM).

Analyzing the latent space of miRNAs frequency table with COMET we identified (Table 1) mir-191 as top marker for cluster 3 (HSC), rank 1, 5 and 205 using respectively CLR, RLE and TMM normalizations, which is associated with the appearance of stem cell-like phenotype in liver epithelial cells [33]. Using TMM normalization for cluster 3 rank 1 is associated with miR-132-3p. miR-187-3p was detected respectively as rank 1 marker for cluster 2 (M) with FQ normalization and as rank 4 in SUM normalization. miR-187 was shown to play a central role in the physiological regulation of IL-10-driven anti-inflammatory responses in TLR4-stimulated monocytes [34].

### SCA analysis of a spatially resolved transcriptomics breast cancer histological section

Spatially resolved transcriptomics (visium technology, 10XGenomics, USA) provides the sequencing of up to 5000 spots (55 µm ∅) of a histological tissue (6.5 × 6.5 mm) section embedded in OTC. This technology does not guarantee single cell sequencing, since in each spot, on the basis of the cells size and density, there could be between 1 and 30 cells.

As further example for the use of SCA we analyzed a breast cancer histological section available as part as the demo data for visium technology. The best clustering could be obtained using SIMLR (Fig. 6A), which generated a partition in 9 clusters where 6 clusters (1, 2, 3, 6, 7, 8) show a very high CSS, Fig. 6B, while other two (5, 8) an intermediate CSS (Fig. 6B). Clusters were then localized on the histological session (Fig. 5C, D). On the basis of an inspection, from two independent pathologists, clusters 1 and 2 (Fig. 6D) were annotated as invasive carcinoma, with a limited similarity with cluster 9 (ductal carcinoma in situ), cluster 5 was assigned to tumor stroma, cluster 7 to invasive carcinoma with undifferentiated/disorganize areas, cluster 6 to invasive carcinoma, showing some morphological characteristics similar to cluster 7 organization. Cluster 8 was assigned to invasive carcinoma, which is however different from the invasive carcinoma observed in cluster 1 and 2. Cluster 9 was assigned to ductal carcinoma in situ. Cluster 3 and 4, because of the limited number of cells could not be assessed by pathologists.

**Fig. 6:**
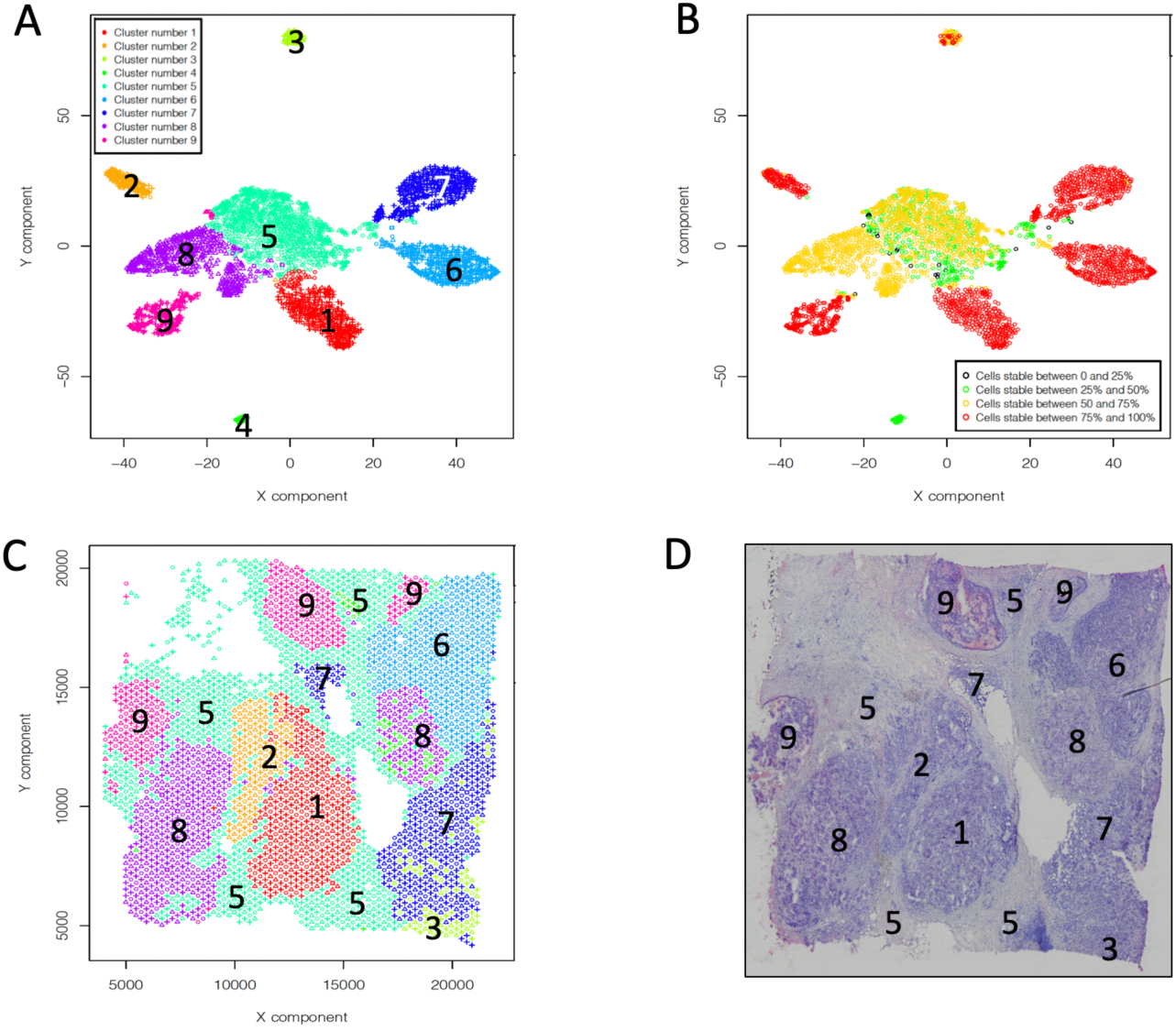
Analysis of human breast cancer (Block A Section 1), from 10XGenomics Visium Spatial Gene Expression 1.0.0. demonstration samples. A) SIMLR partitioning in 9 clusters. B) Cell stability score plot for SIMLR clusters in A. C) SIMLR clusters location in the tissue section. D) Hematoxylin and eosin image.

We tested the ability of SCA to assign TFs to the detected clusters and only cluster 7 could be described by SCA analysis (Fig. 7).

**Fig. 7:**
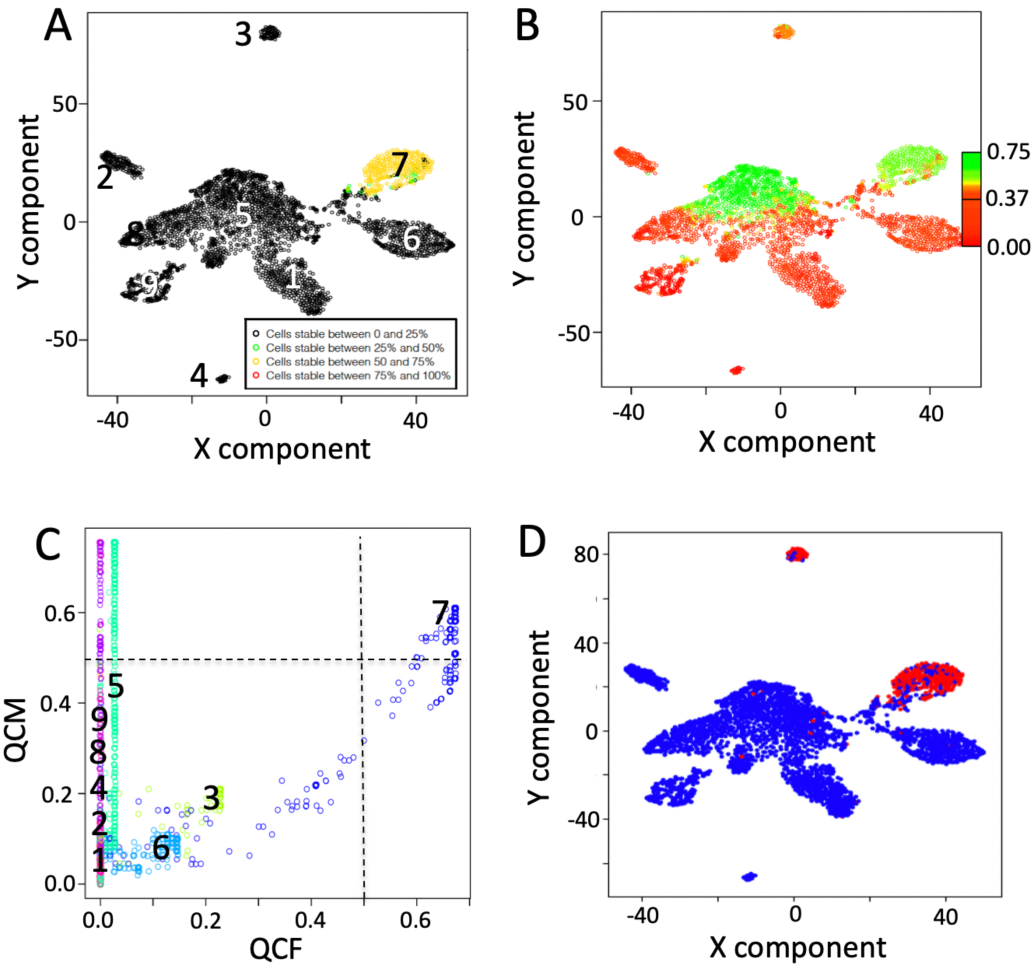
Information contents extracted using as hidden layer the TFs. A) QCF. B) QCM. C) Only cluster 7 show a QCF and a QCF greater than 0.5. D) SOX5 detected as first top ranked gene specific for cluster 7, using as input for COMET the latent space frequency table. Input counts table for SCA are log10 transformed.

COMET analysis of the latent space frequency table provided the detection of SOX5 as top ranked transcription factor specific for cluster 7 (Fig. 7D, Table 2). Notably, SOX5 has been recently associated with breast cancer proliferation and invasion [35], which interestingly correlates with the histological observation that cluster 7 is characterized by a particular undifferentiated structure, which makes it different from the other invasive carcinoma clusters. We also tested immune signature and kinome based SCA, but we could not find any robust association with clustering data (not shown).

**Table 2.**

| *Latent space* | *Cluster* | *Feature 1* | *Feature 2* | *Feature 3* | *Feature 4* | *COMET<sub>sc</sub> statistics* | *TP* | *TN* |
| --- | --- | --- | --- | --- | --- | --- | --- | --- |
| *TFs* | 7 | SOX5 | - | - | - | 2.88E-184 | 0.623 | 0.968 |
| - | 1 | FCGR3B | - | - | - | 1.19E-199 | 0.654 | 0.972 |
| - | 2 | SLC30A8 | - | - | - | 1.67E-67 | 0.194 | 0.992 |
| - | 3 | S100A11 | IFI27 | - | - | 1.86E-89 | 0.670 | 0.997 |
| - | 4 | - | - | - | - | - | - | - |
| - | 5 | S100P<br>Negation | MT2A<br>Negation | PPDPF<br>Negation | CD9<br>Negation | 0 | 0.698 | 0.978 |
|  | 6 | SLITRK6 | - | - | - | 0 | 0.884 | 0.993 |
|  | 7 | SPAG6 | - | - | - | 2.36E-184 | 0.626 | 0.968 |
|  | 8 | - | - | - | - | - | - | - |
|  | 9 | TFF3 | LDHA | - | - | 6.76E-120 | 0.442 | 0.989 |

COMET analysis of the 9 clusters generated by SIMLR using as input the raw counts table, i.e. reference clusters, detected cluster specific markers (Table 2).

We could not find any clear correlation with breast cancer for FCGR3B, SLC30A8, IFI27, SLITRK6, SPAG6. S100A11 is expressed in many cancers [36] and TFF3 was observed to be expressed at higher level in blood of breast cancer patients with metastatic disease with respect to those without [37]. LDHA was shown to act as prognostic biomarker for breast cancer [38].

## Discussion

In this manuscript we highlight that SCA can be uses as tool to uncover hidden features associated to scRNAseq data. Gold and coworkers [8] used SCA as tools for projecting gene-level data onto gene sets. Their observations suggest that SCA may be able to utilize transcription factor target gene sets to help identifying transcription factors with differential activity between conditions or cell types [8]. We decided to used SCA with a different view, we investigated the ability of SCA to reproduce, completely or partially, cell clusters organization depictable from a scRNAseq experiment. Specifically, if SCA encoding is able to reproduce at least one of the clusters, observable aggregating cells on the basis of their full transcriptome, that mean that the encoded biological knowledge is mandatory to obtain a specific aggregation of cells. Here, we show that different hidden layers, derived by experimentally validated data (TFs targets, miRNAs targets, Kinases targets, and cancer-related immune signatures), can be used to grasp single cell cluster-specific functional features. In our implementation, SCA efficacy comes from its ability to reconstruct only specific clusters, thus indicating only those clusters where the SCA encoding space is a key element of a specific cell subgroup. This is clearly demonstrated looking at the top ranked transcription factors, derived from SCA encoded space for cluster 1 and 2 of the blood-cell based dataset (setA), which are key elements for B-cells and monocytes.

A very important element of our SCA implementation is the availability of metrics estimating the robustness of the SCA encoding. CSS [9] is used to evaluate both SCA coherence with respect to full-set based cell clustering (QCF) and SCA model quality (QCM). These metrics are powerful instruments to measure SCA robustness. Furthermore, the effect of SCA input count table normalization on SCA encoding can be estimated using QCF and QCM scores. Thus, allowing to define the optimal condition to retrieve biological knowledge from the SCA encoded space.

Last but not least, since SCA usage could be particularly challenging for life science and medical personnel, lacking of strong computation skill, our implementation of SCA within rCASC framework [9] moderate the above issue, because SCA usage is also fully accessible via GUI.

## Material and Methods

### Datasets

Dataset setA is based on FACS purified cell types [21]. It is made of 100 cells from five cell types (B cells (B), Monocytes (M), Natural Killer (NK), Naïve T cell (N), Hematopoietic stem cells (HSC)). This dataset was analysed without any filtering. Data were log_10_ transformed before clustering.

Dataset HBC_BAS1 is derived from 10XGenomics spatial transcriptomics datasets resources [39]. The filtered sparse matrix from 10XGenomics repository was transformed in a dense matrix using rCASC *h5tocsv* function. Dataset was annotated using ENSEMBL Homo sapiens GRCh38.99 GTF file using the rCASC *scannobyGtf* function. After annotation, ribosomal and mitochondrial protein genes were removed with all ENSEMBL ID not belonging to protein_coding ENSEMBL biotype. Cells with less than 250 detected genes were also removed (i.e. a gene is called detected if it is supported by at least 3 UMIs). After filtering rCASC *topx* function was used to select the 10000 most dispersed genes and from them the 5000 most expressed genes. The final matrix was made by 5000 genes and 3432 cells out of the initial 3813 cells (HBC_BAS1). Data were log_10_ transformed before clustering.

### Model Coding and hyperparameter selection

This work uses sparsely-connected autoencoders [8] (Fig. 1) to grasp cluster-specific hidden features. Autoencoders learning is based on an encoder function that projects input data onto a lower dimensional space. Then, a decoder function recovers the input data from the low-dimensional projections minimising the reconstruction. We implemented the models in python (version 3.7) using TensorFlow package (version 2.0.0), Keras (version 2.3.1), pandas (version 0.25.3), numpy (version 1.17.4), matplotlib (version 3.1.2), sklearn (version 0.22), scipy (version 1.3.3). Optimisation was done using Adam (Adaptive moment estimation) using the following parameters lr=0.01, beta_1=0.9, beta_2=0.999, epsilon=1e-08, decay=0.0, loss=‘mean_squared_error’. RELU (rectified linear unit) was used as activation function for dense layer.

The SCA input counts table is log_10_ transformed or normalized using one of the following tools implemented in rCASC: i) centred log-ratio normalization; ii) relative log-expression; iii) full-quantile normalization; iv) sum scaling normalization; v) weighted trimmed mean of M-values.

### SCA hidden layer definition

SCA hidden layer is generated using as input a tab delimited text file having as first column the feature ids associated to the hidden layer and a second column having the gene associated to a specific feature id. Third column is compulsory and include the reference from which the feature/gene relation was taken.

Experimentally validated transcription factors’ target genes and the associated transcription factor were retrieved from TRRUST v2.0 [12]. Experimentally validated miRNA gene targets and their corresponding miRNA were retrieved from miRTarBase v8.0 [40]. Kinases target genes were retrieved from RegPhos v2.0 database [14]. Cancer immune-signature was manually curated retrieving from PUBMED articles related to the keyword “cancer immune signature” together with genes related to KEGG “Immune system” pathways [41]. Genes associated to KEGG pathways were manually extracted from KEGG pathways public repository [42].

### QCM and QCF metrics

QCM and QCF are extension of CSS [9]. QCF describes the cell stability of a set of reference clusters with respect to clusters generated using SCA latent space information. QCF uses, as CSS, reference clusters, which are those generated using the count table of the full set of cells. These reference clusters can be generated using any of the clustering tools implemented in rCASC [9]. In QCF, reference clusters are compared to multiple runs SCA, where clusters are constructed by the latent space information, using any of the clustering tools implemented in rCASC. High coherence between any of the reference clusters and SCA clusters indicates that the latent space is able to correctly describe at least some the clusters. The QCF threshold for a robust cluster is a value grater of equal to 0.5, i.e. in 50% of the SCA runs cells are colocalizing as in corresponding reference cluster.

QCM is instead measuring the robustness of the SCA model. Since at each run of the SCA the latent space is starting from a random configuration, which is modelled on the basis of the information provided to the SCA, i.e. gene counts, we expect that if the SCA latent space is explaining some of the clusters then those clusters should remain similar between various run of SCA. Thus, QCM measures the reproducibility of each single cluster over a large set of randomly pairs of SCA runs. The lack of reproducibility between clusters over different runs of SCA indicates that latent space information is not related or not robust enough to support conserved cluster structures. The QCM threshold describing a robust model for a cluster is a value grater of equal to 0.5, i.e. in 50% of the SCA runs cells are colocalizing in the same cluster.

### COMET analysis

The cluster-specific markers detection was done using the COMET [20] implementation available in rCASC. COMET was set to extract up to 4 features characterizing each cluster. Although COMET analyses all available clusters, features were investigated only for those clusters characterized by QCM and QCF > 0.5.

### SCA handling functions in rCASC

A full description of the SCA handling functions available in rCASC are available in supplementary data file.

