## supplementary data for "Sparsely-Connected Autoencoder (SCA) for single cell RNAseq data mining"

### Sparsely-Connected Autoencoders (SCAs) for single cell RNAseq data mining supplementary data

21/05/2020

### Contents

### Section 1

A data set, **setA** [*Alessandri et al. 2019*], based on FACS purified cell types [*Zheng et al. 2017*] was used to investigate the SCA behaviour.

#### setA: 100 cells randomly selected for each cell type

- (B) B-cells (25K reads/cell),
- (M) Monocytes (100K reads/cell),
- (HSC) Stem cells (24.7K reads/cell),
- (NK) Natural Killer cells (29K reads/cell),
- (N) Naive T-cells (19K reads/cell)

SetA was previously used to estimate the strength of CSS metric [*Alessandri et al. 2019*]. We clustered setA using all the clustering tools actually implemented in rCASC: tSne+k-mean [*Pezzotti et al. 2017*], SIMLR [*Wang et al. 2017*], griph [*Serra et al. 2019*], Seurat [*Butler et al. 2018*], scanpy [*Wolf et al. 2018*] and SHARP [*Wan et al. 2020*]. All tools but tSne+k-mean and scanpy provided very good and similar partition of the different cell types (Fig. 1).

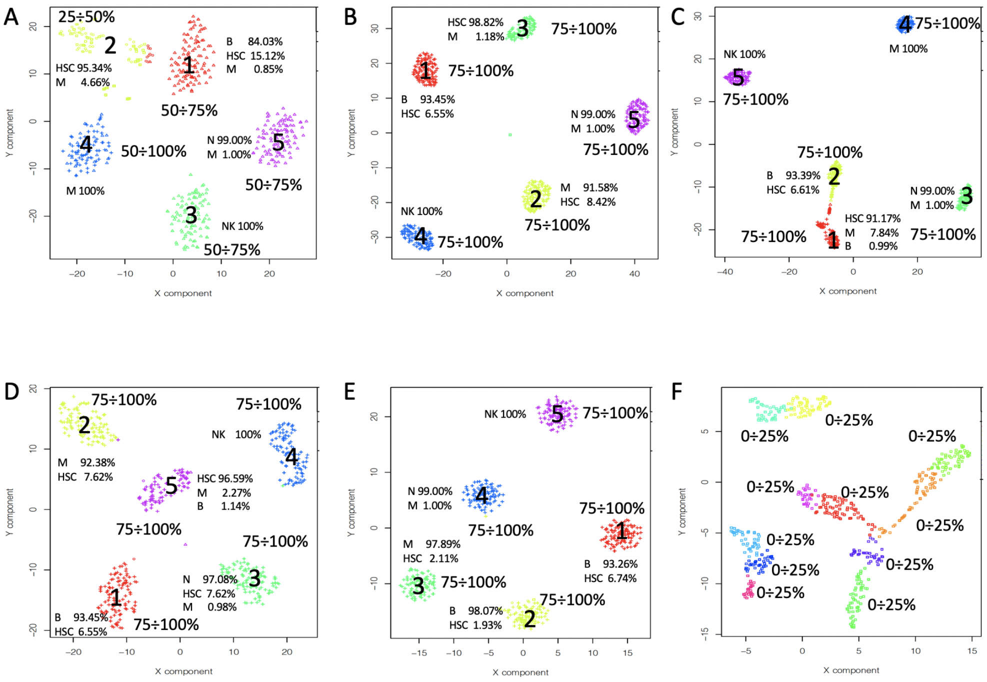

Figure 1: SetA counts table analysis

### Section 2 Mining clusters with Sparsely-Connected Autoencoders (SCAs)

A module for the extraction of cluster-specific functional features was recently implemented in rCASC. This module uses Sparsely-Connected Autoencoders (SCAs) to grasp hidden functional features from the clusters detected using the rCASC clustering tools. The implementation is made for CPU and GPU architectures.

#### Section 2.1: autoencoder function

The function **autoencoder** calculates the neural network.

```
##add example
library(rCASC)
autoencoder(group=c("docker"), scratch=path,
            file=paste(path, "~/newModification/Data/setA.csv", sep="/"),
            separator=",", nCluster=5, bias="kinasi", permutation=160,
            nEpochs=2000, patiencePercentage=5,
            cl=paste(path,"~/newModification/Data/Results/setA/5/setA_clustering.output.csv",sep=""),
```

```
          seed=1111, projectName="setAKinasi", bN="NULL")
```

- *autoencoder parameters* (only those without default; for the full list of parameters please refer to the function help):
  - *bias* ("mirna" , "TF", "CUSTOM", "kinasi","immunoSignature") refers to the rules used to generate the partially connected hidden layer. If CUSTOM is selected the path to the file required to build the autoencoder structure should be assigned to the parameter bN. The hidden layer description file must have the following structure source (header is mandatory and row names must not be present, any extra column will be not taken in account):
    * name of the hidden layer node, column 1
    * geneTarget, i.e. genes present in the hidden node, column 2.
  - *nEpochs* is the parameter referring to the number of times the training vectors are used to update the weights. At the end of this learning step the best weights for nodes will be defined. Please note that not necessarily the best weights are in the last epoch. Frequently, they are located in the first 1000 epochs. Thus, a good approach is to run 2000 epochs with 1 permutation only and check the learning rate, Figure 2. The optimal epochs is selected within the flat part of the learning curve. Then, the analysis is run again with the number of permutations of your choice.
  - *permutation* refers to the number of times the neural network is calculated from scratch. The number of permutations depends on the level of consistency requested, usually 160 permutations provides robust results, but it might take some times to execute on a common laptop , thus lower number of permutations might be selected, e.g. 80.
  - *cl* refers to the full path to the **_clustering.output** file to be used, which was generated using rCASC clustering tools.
  - *ProjectName* parameter refers to the name of the folder where **autoencoder** outputs will be saved.

The **autoencoder** function produces as output many files with the extension **Xdensespace.format**, in the folder **ProjectName/Permutation**. The number of such files corresponds to the number of selected permutations.

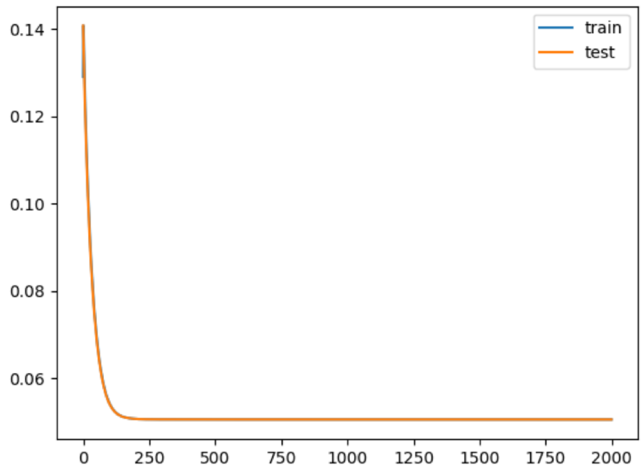

Figure 2: Learning rate saved in learning.jpg. In this example the maximum level of learning is obtained with 250 epochs. Thus the analysis with multiple permutations can be done setting epoch slightly greater than 250, e.g. 500

##### Section 2.2: autoencoderClustering function

The function **autoencoderClustering** generates the numerical data required to understand if there is coherence between counts table derived clusters and those derived using autoencoders latent space.

```
##add example
library(rCASC)
autoencoderClustering(group=c("docker"), scratch.folder=path,
                      file="~/newModification/Data/Results/setAKinasi/setA.csv",
                      separator=",", nCluster=5, clusterMethod="SIMLR", seed=1111,
                      projectName="setAKinasi",pcaDimensions=15,permAtTime=4,
                      largeScale=TRUE)
```

- *autoencoderClustering parameters* (only those without default; for the full list of parameters please refer to the function help):
  - **IMPORTANT**: *file* parameter refers to the full path of the count table copied in the results output folder of the **autoencoder** function.
  - *projectName* parameter is the same indicated in **autoencoder** function.
  - *clusterMethod* parameter refers to any of the clustering methods available in rCASC

(“GRIPH”,“SIMLR”,“SEURAT”,“SHARP”), not necessary should be the same used for the raw count table clustering, actually it is useful to test more than one clustering method to see which one provides the best convergence with the clusters initially generated using the genes counts table.

– *pcaDimensions* parameter is only required when Seurat is used as clustering tool.

The **autoencoderClustering** output is a folder where the name is given by the combination of the *projectName* and the *clusterMethod*, e.g in the above example **setAKinasi__SIMLR**. In this folder the is the file **label.csv**, where is located the cluster assignment for each cells over each permutation.

The output of **autoencoderClustering** is used by the function **autoencoderAnalysis** to generate the QCF and QCM statistics (Figure 3). QCF is an extension of CSS and it measures the ability of latent space to keep aggregated cells belonging to predefined clusters generated using the gene count table. The metrics has the range between 0 and 1, where 1 indicates a high coexistence of cells within the same cluster in the analysis, and 0 a total lack of coexistence of cells within the same cluster. In the figure 3A, only cluster 7 can be well explained by latent space, 3B. QCM metric is also an extension of CSS and it measures the ability of the neural network to generate consistent data over the different training. In the Figure 3C, SCA provides consistent data only for clusters 5 and 7. Informative clusters are those characterised by high QCM and QCF scores, Figure 3D only cluster 7 is characterized by a robust neural network able to keep the cell aggregated using hidden layer knowledge. Dashed red line (Figure 3D) indicates the defined threshold to consider the latent space information suitable to support cells’ clusters.

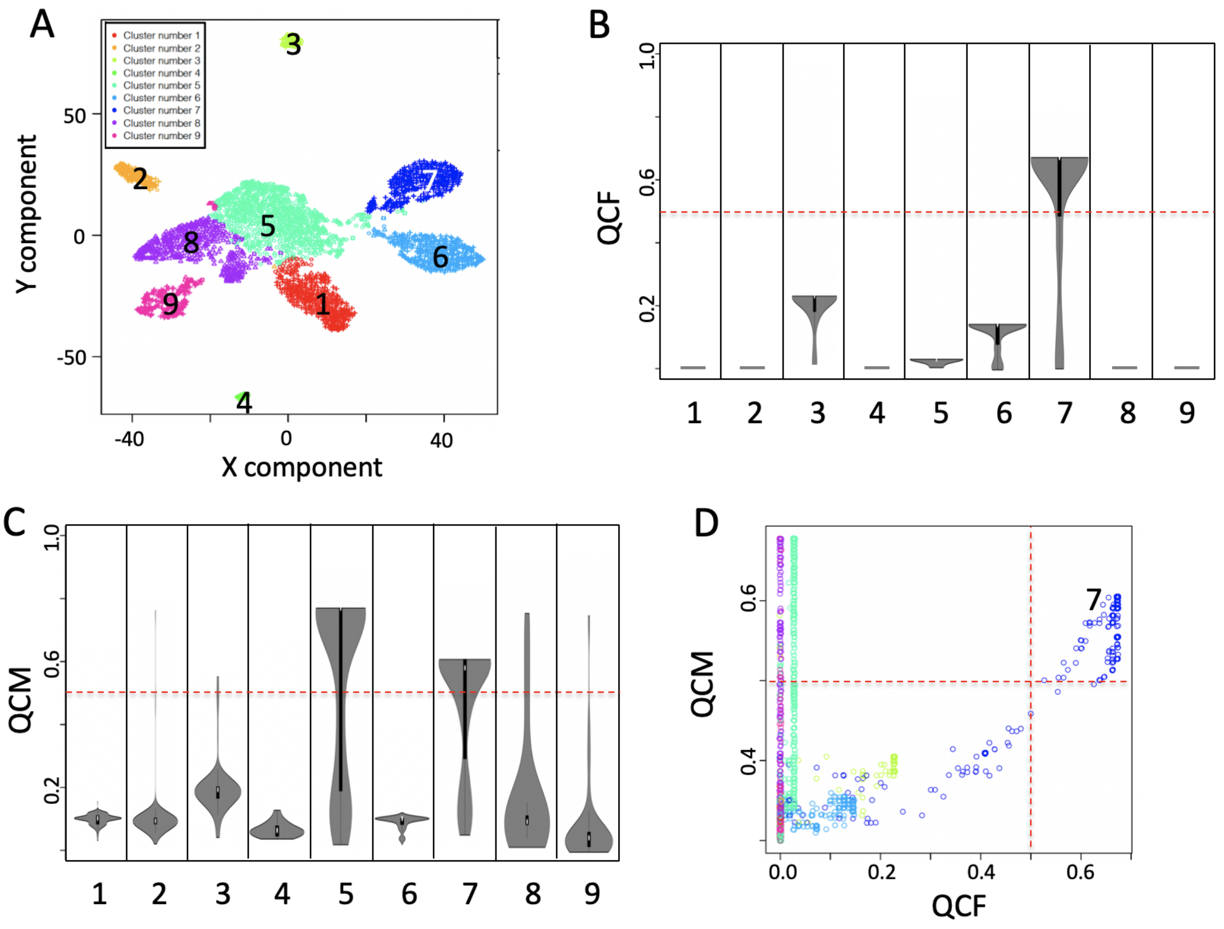

Figure 3: Breast cancer data set in Fig. 5 main manuscript. A) Nine clusters were detected analysing breast cancer dataset with SIMLR. B) QCF violin plot. C) QCM violin plot, D) Combined view of QCM and QCF

**Section 2.3: autoencoderAnalysis function**

Function **autoencoderAnalysis** transforms **autoencoderClustering** function outputs in readable results.

```
library(rCASC)
autoencoderAnalysis(group=c("docker"), scratch.folder=path,
                    file="~/newModification/Data/Results/setAKinasi_SIMLR/setA.csv",
                    separator=",", nCluster=5, seed=1111,
                    projectName="setAKinasi_SIMLR", Sp=0.8)
```

- *autoencoderAnalysis parameters* (only those without default; for the full list of parameters please refer to the function help):
  - file e projectName parameters refer to the outputs of **autoencoderClustering**

  - *sp* is the similarity threshold defined for CSS (please see *Section 5.1 Cell Stability Score: mathematical description in Alessandri et al. 2019 supplementary data*). This paramenter has as default 0.8. Reduction below 0.7 of this threshold reduces the specificity of CSS; values equal or smaller than 0.5 are meaningless.

The outputs of **autoencoderAnalysis** are the pdfs:

- **setA__stabilityPlot.pdf** (QCF, Figure 3B)
- **setA__stabilityPlotUNBIAS.pdf** (QCM, Figure 3C)
- **setA__StabilitySignificativityJittered.pdf** (QCM versus QCF, Figure 3D)

###### Section 2.4: autoFeature function

The **autoFeature** creates the frequency table for COMET analysis.

```
library(rCASC)
autoFeature(group=c("docker"), scratch.folder=path,
            file="~/newModification/Data/Results/setAKinasi_SIMLR/setA.csv",
            separator=",", nCluster=5,projectName="setAKinasi_SIMLR")
```

**IMPORTANT**: this function produces an output for all clusters, but only results related to clusters supported by QCM and QCF means greater than 0.5 have to be taken in account.

- *autoFeature parameters* (only those without default; for the full list of parameters please refer to the function help):
  - file e projectName parameters refer to the outputs of **autoencoderAnalysis**

###### Section 2.5: cometsc2 function

**cometsc2** is a modification of **cometsc** function suitable to handle **autoencoder** frequency table, i.e. **autoencoder** frequency table contains the % of permutations in which each hidden node for each cell was characterized by a weight different from 0 (if a hidden node is characterized by a weight equal to 0 it means that it was not used in that specific permutation). **autoFeature** function is used to extract functional features from autoencoders latent space frequency table output.

```
library(rCASC)
cometsc2(group=c("docker"),
         file="~/newModification/Data/Results/setAKinasi_SIMLR/5/freqMatrix.csv",
         scratch.folder=path, threads=1, X=0.15, K=2, counts=c("True", "False"),
         skipvis=c("True", "False"), nCluster=5, separator=",",
         clustering.output=paste(path,"~/newModification/Data/Results/setA/5/setA_clustering.output.csv
                                 sep=""))
```

- *cometsc2 parameters* (only those without default; for the full list of parameters please refer to the function help):
  - *file* is the path to the frequency matrix generated by **autoFeature** function.
  - *clustering.output* is the path to the clustering.output generated by rCASC on the initial counts table

**cometsc2** outputs are the same of COMET, for more information please refer to *COMET documentation*.

**IMPORTANT**: this function produces an output including all clusters, but **only results related to clusters supported by QCM and QCF means greater than 0.5 have to be taken in account**.

##### Section 2.6: wrapperAutoencoder function

The function **wrapperAutoencoder** executes all the steps required for the autoencoder analysis. It should be used when it is clear the overall set of steps to be performed and a set of analyses with the same parameters are done.

```
library(rCASC)
wrapperAutoencoder(group="docker", scratch.folder=scratch.folder,
                   file="/home/lucastormreig/test/setA.csv", separator=",",
                   nCluster=5, bias="mirna", permutation=10, nEpochs=10,
                   cl="/home/lucastormreig/test/setA_clustering.output.csv",
                   projectName="mirna",clusterMethod="GRIPH")
```

- *wrapperAutoencoder parameters*:
  - *group*, a character string. Two options: sudo or docker, depending to which group the user belongs
  - *scratch.folder*, a character string indicating the path of the scratch folder

- *file*, a character string indicating the path of the file, with file name and extension included
- *separator*, separator used in count file, e.g. '\t', ','
- *nCluster*, number of cluster in which the dataset is divided
- *bias*, bias method to use : "mirna" , "TF", "CUSTOM", kinasi,immunoSignature
- *permutation*, number of permutations to perform the pValue to evaluate clustering
- *nEpochs*, number of Epochs for neural network training
- *patiencePercentage*, number of Epochs percentage of not training before to stop.
- *cl*, Clustering.output file. Can be the output of every clustering algorithm from rCASC or can be customized with first column cells names, second column cluster they belong.All path needs to be provided.
- *seed*, important value to reproduce the same results with same input
- *projectName*, might be different from the matrixname in order to perform different analysis on the same dataset
- *bN*, name of the custom bias file. This file need header, in the first column has to be the source and in the second column the gene symbol.All path needs to be provided,
- *lr*, learning rate, the speed of learning. Higher value may increase the speed of convergence but may also be not very precise
- *beta_1*, look at keras optimizer parameters
- *beta_2*, look at keras optimizer parameters
- *epsilon*, look at keras optimizer parameters
- *decay*, look at keras optimizer parameters
- *loss*, loss of function to use, for other loss of function check the keras loss of functions.
- *clusterMethod*, clustering methods: "GRIPH","SIMLR","SEURAT","SHARP"
- *pcaDimensions*, number of dimensions to use for Seurat Pca reduction.
- *permAtTime*, number of permutation in parallel
- *largeScale*, boolean for SIMLR analysis, TRUE if rows are less then columns or if the computational time are huge

- *Sp*, minimun number of percentage of cells that has to be in common between two permutation to be the same cluster.
- *threads*, integer refering to the max number of process run in parallel default 1 max the number of clusters under analysis, i.e. nCluster
- *X*, from 0 to 1 argument for XL-mHG default 0.15, for more info see cometsc help.
- *K*, the number of gene combinations to be considered., possible values 2, 3, 4, default 2. WARNING increasing the number of combinations makes the matrices very big
- *counts*, if set to True it will graph the log(expression+1). To be used if unlogged data are provided
- *skipvis*, set to True to skip visualizations
